# Vegetative phase change in *Populus tremula x alba*

**DOI:** 10.1101/2020.06.21.163469

**Authors:** Erica H. Lawrence, Aaron R. Leichty, Erin E. Doody, Cathleen Ma, Steven H. Strauss, R. Scott Poethig

## Abstract

- Plants transition through juvenile and adult phases of vegetative development in a process known as vegetative phase change (VPC). In poplars (genus *Populus*) the differences between these stages are subtle, making it difficult to determine when this transition occurs. Previous studies of VPC in poplars have relied on plants propagated *in vitro*, leaving the natural progression of this process unknown.
- We examined developmental morphology of seed-grown and *in vitro* derived *Populus tremula x alba* (clone 717-1B4), and compared the phenotype of these, to transgenics with manipulated miR156 expression, the master regulator of VPC.
- In seed-grown plants, most traits changed from node-to-node during the first 3 months of development but remained constant after node 25. Many traits remained unchanged in clones over-expressing miR156, or were enhanced when miR156 was lowered, demonstrating their natural progression is regulated by the miR156/SPL pathway. The characteristic leaf fluttering of *Populus* is one of these miR156-regulated traits.
- Vegetative development in plants grown from culture mirrored that of seed-grown plants, allowing direct comparison between plants often used in research and those found in nature. These results provide a foundation for further research on the role of VPC in the ecology and evolution of this economically important genus.

## Introduction

During ontogeny, plants transition through distinct phases, which is reflected in morphological variation in the organs produced across development. These developmental transitions include the visually obvious transition between vegetative and reproductive growth as well as the often more subtle change between juvenile and adult vegetative stages, known as vegetative phase change (Ashby, 1948; Brink, 1962; Doorenbos, 1965; Poethig, 2013). In species with abrupt changes in vegetative morphology—termed *heteroblastic*—the distinction between the juvenile and adult phases is unambiguous (Zotz *et al*., 2011). For example, many species of *Acacia* transition from producing bipinnate compound leaves to producing a simple undivided leaf termed a phyllode (Walters & Bartholomew, 1984). Other examples include *Eucalyptus globulus*, which transitions from producing ovate, wax-covered leaves to producing lanceolate glaucous leaves (James & Bell, 2001) and *Ipomoea caerulea*, which transitions from producing monolobed to trilobed leaves (Ashby, 1948). Vegetative phase change is more difficult to identify in species that experience less drastic changes in leaf and shoot morphology. As a result, we know little about the duration of juvenile and adult phases in different taxa, or the relationship between this developmental transition and other aspects of plant life history, such as plant responses to biotic and abiotic conditions. However, even in these so-called *homoblastic* species (Goebel, 1900), organs produced early in shoot development differ from those produced later in development in a variety of morphological, anatomical, and physiological traits (Feng *et al*., 2016). Identifying these traits and determining which are components of vegetative phase change as opposed to those regulated by other factors, such as plant size or chronological age, is an important problem in plant development.

In *Arabidopsis* and most other plants, the transition from the juvenile to the adult phase of vegetative development is regulated by the closely related microRNAs, miR156 and miR157 (Poethig, 2013; He *et al*., 2018). These miRNAs directly repress members of the Squamosa Promoter Binding Protein-Like (*SPL*) family of transcription factors (Rhoades *et al*., 2002; Schwab *et al*., 2005) which promote adult traits (Xu *et al*., 2016). During the juvenile phase miR156/157 levels are high, but as plants age their expression declines, allowing for the increased expression of their *SPL* targets. This coordinated change in miR156 and *SPL* gene expression leads to the onset of adult traits, with different traits typically appearing at slightly different times in shoot development, depending on their sensitivity to the level of SPL proteins (He *et al*., 2018).

Vegetative phase change is likely to be of particular importance in long-lived perennial species such as trees, because the ability of these species to survive to reproductive competence requires that they cope with changing conditions over a long period of time. Poplars are of considerable economic and ecological importance and have become popular model systems for tree biology due to their ability to be propagated in culture and susceptibility to *Agrobacterium*-mediated transformation (Taylor, 2002). Leaf and shoot phenotypes of plants derived from culture and over-expressing miR156 have been described in clones of *Populus x canendensis* (Wang *et al*., 2011) and *P. tremula x alba* (Rubinelli *et al*., 2013). However, the significance of this information for vegetative phase change in *Populus* is still unknown because these studies lacked the earliest developmental stages and fine-scale timeline needed to capture vegetative phase change, and relied on gain-of-function transgenics. Here, we compare the vegetative phenotype of seed-grown *Populus tremula x alba* with plants originating from explants in tissue culture across multiple timepoints during vegetative development, and identify phase-specific traits using both gain- and loss-of-function miR156 transgenics. We found that shoot development in plants propagated *in vitro* closely resembles the development of seed-grown plants, and that vegetative phase change occurs very early in both types of shoots. Most of the traits we examined changed significantly across shoot development and were sensitive to the level of miR156. One of these was leaf fluttering, a trait that has a significant effect on photosynthetic efficiency (Roden & Pearcy, 1993b). Our results provide basic information about the nature of vegetative phase change in *Populus tremula x alba* and demonstrate that shoots regenerated *in vitro* can be used as models of primary shoot development in this species.

## Materials and Methods

### Plant transformation

Assembly of Ubi10:AthMIR156a and Ubi10:MIM156 constructs have previously been described (Feng *et al*., 2016). Additionally, a second MIM156 construct was designed with Golden Gate Cloning (Engler *et al*., 2014) and used for a later transformation after losing many of the original lines to a chamber malfunction. Further details of construct assembly can be found in Table S1.

Poplar line 717-1B4 was transformed using *Agrobacterium tumefaciences* (strain *AGL1*) following the protocol described in Filichkin *et al*. (2006). Briefly, leaf and stem explants were selected on callus-induction, shoot induction, shoot elongation and rooting media using BASTA. A total of 30 Ubi10:AthMIR156a events and 25 Ubi10:MIM156 events were isolated. PCR was used to first screen for transgene insertion using two primer sets, each utilizing a primer to the *Arabidopsis* UBIQUITIN 10 promoter and either the *MIR156a* or MIM156 sequence (Table S2). Of these, we screened 28 Ubi:AthMIR156a lines and 19 Ubi10:MIM156 lines for levels of miR156 and miR157 (Fig. S**1A-B**). Based on levels of miR156, two lines for each construct type were selected (or based on availability in the case of event GG.MIM156-22). For the Ubi10:AthMIR156a and Ubi10:MIM156 constructs made with the pCambia 3300 backbone, the Ubi10:AthMIR156a construct had a higher rate of producing transgenic shoots (percent of explants forming shoots was 52% versus 33.9%). We did not track efficiency for the other MIM156 construct.

### Plant material and growth conditions

*Populus tremula x alba* line 717-1B4, two independent miR156 over-expressing (miR156OX) lines, 40 and 78, and two independent MIM156 lines, 22 and 84, were obtained by *in vitro* propagation and hardening on propagation media as described in Meilan and Ma (2006). Plants were then transplanted to Fafard-2 growing mix (Sungro Horticulture, Massachusetts, USA) in 0.3-L pots in the greenhouse at the University of Pennsylvania (39.9493°N, 75.1995°W, 22.38 m a.s.l.) and kept in plastic bags for increased humidity for 2 weeks. Plants were transferred to 4.2-L pots with Fafard-52 growing mix 3 weeks later and fertilized with Osmocote 14-14-14 (The Scotts Company, Marysville, OH, USA), supplemented by weekly applications of Peters 20-10-20 (ICL Fertilizers, Dublin, OH, USA). Plants were grown at a temperature ranging from 22 and 27°C on a 16 hr photoperiod, with illumination provided by a combination of natural light and 400-W metal halide lamps (P.L. Light Systems, Ontario, Canada) with daily irradiances between 300 and 1,500 μmol m^-2^ s^-1^ measured at the top of the canopy across the day. Planting of individuals was staggered across three months with the position of plants in the greenhouse randomized and individuals rotated frequently.

*Populus tremula x alba* seeds were purchased from Sheffield’s Seed Company (Locke, NY), and were germinated on a layer of vermiculite on top of Fafard-2 growing mix in 0.64-L pots in the greenhouse under conditions described above. Seedlings were transplanted one month after germination into Fafard-52 growing mix in 1.76-L pots, fertilized with Osmocote 14-14-14. They were transplanted to 4.2-L pots 3 months later. Planting of individuals was staggered across five months and individuals were rotated within the greenhouse frequently.

### mRNA and smRNA abundance

For initial screening of miR156 and miR157 abundance, and follow-up validation of selected lines, shoot apices with leaf primordia less than 3 cm were used. For characterization of temporal patterns of miR156/157 and *SPL* abundance in seed-grown and culture derived plants, leaf primordia smaller than 1 cm were sampled and pooled with at least 3 leaves per replicate. For all plants, node or leaf positions are counted from the base of the plant where the first leaf produced is number 1. Primordia were collected across the plant’s lifespan to obtain samples for each of the measured leaf positions when they were at the same stage of development, frozen with liquid N_2_, and stored at −70°C until all samples were ready for RNA extraction. Total RNA was isolated using the Spectrum™ Plant Total RNA Kit (Sigma-Aldrich), according to the manufacturer’s instructions. smRNA levels were measured by RT-qPCR, using primers specific for mature sequences of miR156, miR157 and miR159 in combination with the stem-loop RT primers described in Varkonyi-Gasic et al. (2007) (Table S2). miR159 was used as an endogenous control (Leichty & Poethig, 2019) and validated across time points in seed-grown and culture derived plants (Fig. **S2**). cDNA was synthesized using Invitrogen SuperScript III following the methods of Varkonyi-Gasic et al. (2007). Platinum Taq (Invitrogen) was used with the Roche universal hydrolysis probe #21, and a three-step amplification protocol. For measuring mRNA abundance, total RNA was DNase digested (Qiagen) followed by cDNA synthesis using Invitrogen SuperScript III using each manufacturer’s protocol and a polyT primer. *PtaACT2* or *PtaCDC2* was used as the endogenous control (Pettengill *et al*., 2012). Resulting cDNA was quantified by qPCR using SYBR-Green Master Mix (Bimake) and primers specific to the endogenous control and *SPL* genes (Table S2) with a three-step amplification protocol. Relative measures of abundance were calculated using the 2^-ΔΔCt^ method (Livak & Schmittgen, 2001).

### Stomatal impressions, epidermal traits, petiole morphology and abaxial trichome density

Stomatal density and epidermal cell size were measured using clear nail polish impressions of adaxial and abaxial surfaces of fully expanded leaves. Trichomes were removed from the abaxial side of leaves with tape when necessary, and three sections per leaf were imaged and photographed using an Olympus BX51 light microscope. Hand sections of leaf petioles within 1 cm of leaf attachment site were imaged as described above. Trichome density on the abaxial side of the leaf was measured using an Olympus MVX10 dissecting scope. Measurements were made along the mid vein, where the density was low enough to permit accurate counts. All measurements and counts were made using Fiji software (Schindelin *et al*., 2012).

### Leaf vein densities

Sections from new fully expanded leaves were cleared using NaOH and bleach, and then stained with Safranin following the protocol described in Scoffoni and Sack (2013). Vein lengths of all visible vein orders were measured from microscope images using Fiji software (Schindelin *et al*., 2012), and vein densities were calculated following Scoffoni and Sack (2013).

### Leaf shape and size

Fully expanded leaves were attached to paper using double-sided tape, labeled by node number increasing from the base of the plant, and scanned. The scanned images were measured using the Morphological Analysis of Size and Shape (MASS) software (Chuanromanee *et al*., 2019). The SHAPE program (Iwata & Ukai, 2002) and Fiji (Schindelin *et al*., 2012) was used to determine the number of serrations and the perimeter of the leaf. The perimeter of the leaf was measured from the tops of the serrations so as not to include the added length from the serration indentations. Serrations were analyzed relative to the leaf perimeter so that the data were not influenced by differences in overall leaf size.

### Leaf initiation rate, internode distance and branching

The leaf initiation rate was determined by counting the node of the uppermost visible leaf at least twice a week between 10 and 160 days of growth in seed-grown plants and between 32 and 55 days after transplant to soil in 717-1B4 lines. The number of branches at least 1 cm in length between nodes 1 and 25 was determined after 4 months of growth in soil.

### Fluttering

To measure leaf fluttering, plants were placed 100 cm away from a fan, with the leaf to be measured facing perpendicular to the wind. Wind speeds at the sample leaf position were 1.5 m/s as measured with an anemometer. Leaf movement was filmed at 240 fps for 30 seconds (representative example in supplemental file “Poplar Fluttering Video”). Leaf movement was analyzed by following the movement of a mark placed at the tip of the leaf using Tracker video analysis and modeling tool v5.0.7. The height and width of the petiole were measured using a digital caliper.

### Statistical Analysis

All statistical analyses were performed in JMP ^®^ Pro v. 14.0.0 (SAS Institute Inc., Cary, NC). Traits in seed-grown poplar across leaf positions were measured from a minimum of 42 individuals and fit with a linear or second-degree polynomial as appropriate with *p*-values and *R*^2^ values reported. Traits that were examined at multiple leaf positions in 717 wildtype, MIM156 lines 84 and 22 and miR156OX lines 40 and 78 were measured from a minimum of 6 individuals and compared by ANCOVA and a Student’s *t* test (*α* = 0.05) where leaf position and genotype were the main effects. Whole-plant measures of branching and internode length in these plants was compared by one-way ANOVA where genotype was the main effect and a Student’s *t* test (*α* = 0.05), and leaf emergence was compared by ANCOVA with days after transplant and genotype as the main effects.

## Results

### Isolation of MIR156a over-expressing and MIM156 plants

For the lines with confirmed transgene insertions, the relative abundance of miR156 averaged 0.63 and 1.37 for the MIM156 and *MIR156a* lines, respectively (Figure **S1A**,**B**). Lines MIR156A-78 and MIR156-40 were selected as strong and moderate miR156 over-expressors, respectively (Figure **S1A**). Line MIM156-84 was selected as a strong miR156 mimic, while line MIM156-22 was selected to represent a transgenic line in which the miR156 level was not significantly different from wildtype (Figure **S1A**,**B**, *T*-test: *p* = 0.51). The levels of miR156-targeted *SPL* genes was consistent with the levels of miR156/157 in different transgenic lines. MIM156-84 had, respectively, 3- and 9-fold more *PtSPL24* and *PtSPL23* transcripts relative to wildtype (Figure **S1C**). As expected based on its levels of miR156, the transcripts of both *PtSPL24* and *PtSPL23* in MIM156-22 did not differ from wildtype (Figure **S1C**). Both MIR156A-40 and MIR156A-78 showed significantly lower levels of these same genes when compared with wildtype (Figure **S1C**). To confirm the specificity of these transgenes, we also measured levels of *PtSPL4* and *PtSPL7*—genes with no predicted miR156 target site. In both cases, there was no clear pattern of transcript abundance across transgenic lines (Figure **S1C**).

### miR156/157 and ptSPL24 abundance in seed-grown plants and 717-1B4 lines

Patterns of miR156/157 abundance in the leaf primordia of seed-grown plants matched that previously reported for other species, such as *A. thaliana* and *Vachellia* sp. (He *et al*., 2018; Leichty & Poethig, 2019). miR156/157 abundance dropped 7-fold between nodes 1 and 10, followed by a slower decline in successive nodes (Fig. **1A**). We first determined that the control small RNA, miR159, was present at the same level in leaves 15 and 25 of seed-grown plants and the clone 717-1B4 (Fig. **S2**), and then used this transcript to normalize the level of miR156/157 in wild-type and transgenic genotypes of 717-1B4. The abundance of miR156/157 in the primordia of leaf 15 and 25 in wildtype 717-1B4 was similar to the amount of miR156/157 in these same leaves in seed-grown poplar (Fig. **1A, B**), suggesting that miR156/157 are expressed at comparable levels at the same node in plants propagated by these methods. Leaf 15 and 25 of the miR156OX lines had, respectively 4.9 and 5.6 times the amount of miR156 as wild-type leaves, a level comparable to that of leaf 3 and 5 of seed-grown plants (Fig. **1A, B**). Although the amount of miR156/157 in leaves 15 and 25 of the MIM156 lines was similar to that in 717-1B4 and seed-grown poplar (Fig. **1A, B**), line 84 had approximately 1/3 the amount of miR156 at an earlier stage of development, before it was transferred to soil (Fig. **S1A**). Target mimics do not necessarily reduce the level of their target miRNAs (Franco-Zorrilla *et al*., 2007), so the significance of these MIM156 data is difficult to interpret. In any case, the similarity between the amount of miR156/157 in MIM156 and wild-type lines shows that the transformation protocol and vector components have no independent effect on miR156/157 abundance.

**Figure 1.**
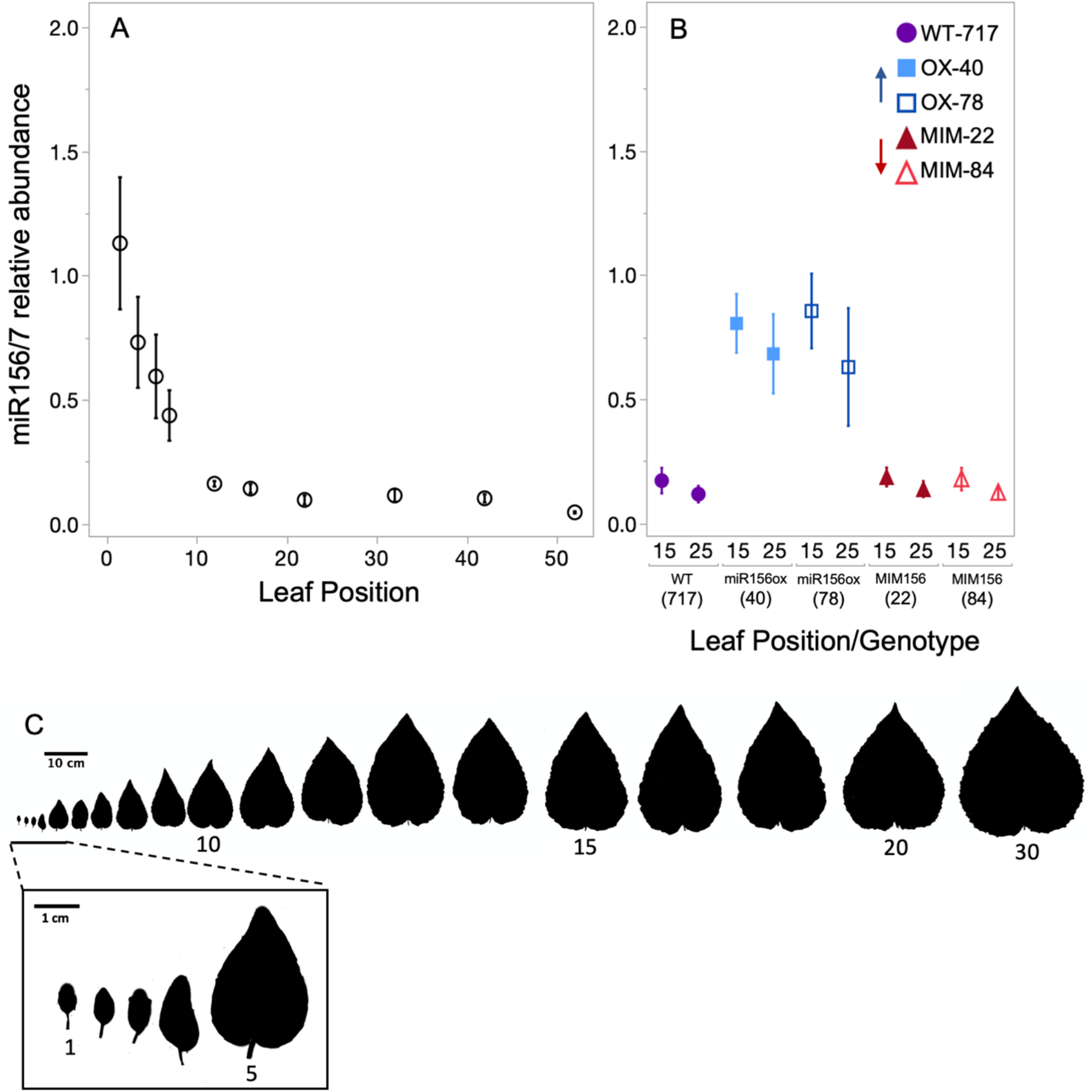
The relative abundance of miR156/157 in *Populus tremula x alba* leaf primordia grown from seed (A) or in transgenic and wildtype plants of the 717-1B4 clone initially propagated in tissue culture (B). All miR156/157 measurements are relative to the average of 4 replicates from seed-grown leaves 1 & 2. Data are presented as means ± s.e.m (n = 3) with seed-grown data depicted as black open circles, 717 wildtype by purple circles, miR156a overexpression lines depicted by blue squares and MIM156 lines depicted by red triangles. (C) Scans of successive leaves from nodes 1-17, 20 and 30 from plants grown from seed.

To determine if the changes in miR156/157 abundance observed in seed- and culture-derived plants were large enough to have downstream effects on the expression of their *SPL* targets, we measured *ptSPL24* abundance across nodes 1 and 34 in seed-grown plants and at nodes 15 and 25 of 717-1B4 lines. *ptSPL24* abundance increased more than 11-fold between nodes 1 and 34 in seed-grown plants (Fig. **2A**). In wildtype 717-1B4, *ptSPL24* abundance increased 2-fold between nodes 15 and 25 (Fig. **2B**). Across a similar span in seed-grown plants, *ptSPL24* abundance increased by more than 1.5-fold between nodes 12 and 22, again indicating that the expression patterns of genes in the miR156/157-SPL pathway are comparable between similar nodes of plants propagated via seeds and cuttings. Leaves 15 and 25 in the miR156OX lines had 4- and 15-fold less *ptSPL24* respectively, compared to these same nodes in the wildtype line. This is consistent with the levels of miR156/157 in these lines, and indicates that these levels of miR156/157 are functionally significant. The abundance of *ptSPL24* in MIM156 line 22 was similar to wild-type, consistent with the observation that the abundance of miR156 in this line is not significantly different from wildtype. Interestingly, the abundance of *ptSPL24* was more than 2-fold greater in leaf 15 of MIM156 line 84 compared to wildtype, although the level of miR156/157 in this transgenic leaf is similar to wildtype. There was no difference in the abundance of *ptSPL24* at node 25 in MIM156 line 8 and wildtype plants . Together with the abundance of miR156 and *ptSPL24* measured in young plants still in culture (Fig. **S1**), these data suggest that the MIM156 transgene has the greatest effect during the juvenile phase, when miR156 levels are naturally high.

**Figure 2.**
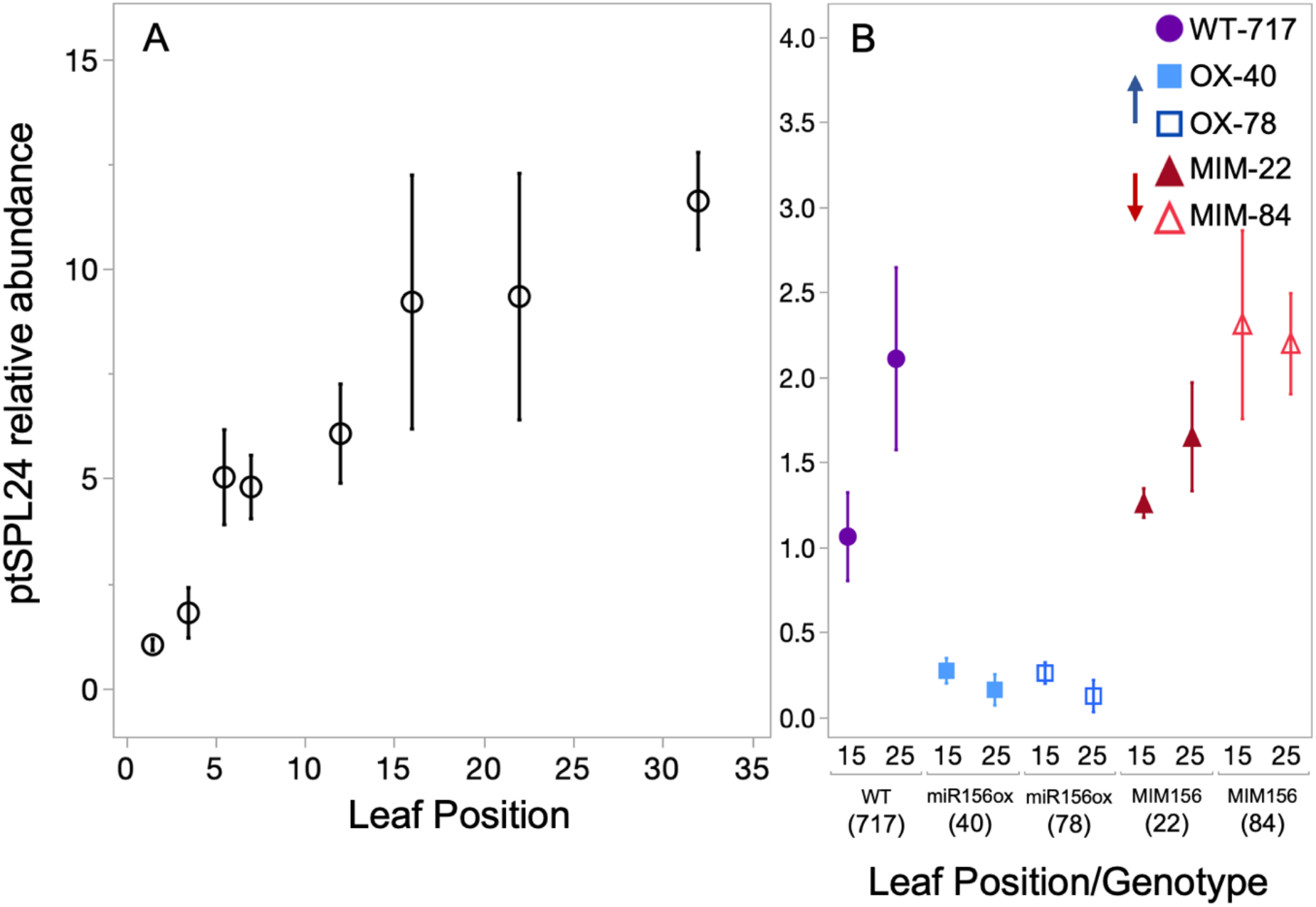
The relative abundance of *ptSPL24* transcripts in *Populus tremula x alba* leaf primordia grown from seed (A) or in transgenic and wildtype plants of the 717-1B4 background initially propagated in tissue culture (B). The abundance of *ptSPL24* in seed-grown plants relative to seed-grown leaves 1 & 2 (A) or relative to the abundance in wildtype 717 leaf 15 in plants initially propagated in tissue culture (B). Data presented as means ± s.e.m (n = 3) with see-grown data depicted as black open circles, 717 wildtype by purple circles, miR156a overexpression lines depicted by blue squares and MIM156 lines depicted by red triangles.

### Changes in miR156 abundance alter plant growth rate and form

Plants grown from seed produced leaves at a relatively slow rate for the first few weeks after planting, and transitioned to a faster, linear growth rate after producing 15-20 leaves (Fig. **3A**). Leaf initiation in wild-type and transgenic clones of 717-1B4 was delayed for 35-40 days after transplanting to soil, and then occurred at increasing rate over the next 2-3 weeks of measurement. During this latter period, miR156OX lines had higher rates of leaf initiation than either wild-type or MIM156 lines (Fig. **3B**). miR156OX lines also had shorter internodes than either the wild-type and MIM156 lines (Fig. **3C**) and produced significantly more axillary branches after 3 months than these genotypes (Fig. **3D**, Photographs of plants at this age in Fig. **S3**). The MIM156 lines were not significantly different from 717-1B4 for any of these traits, although these lines did have slightly longer internodes than 717-1B4.

**Figure 3.**
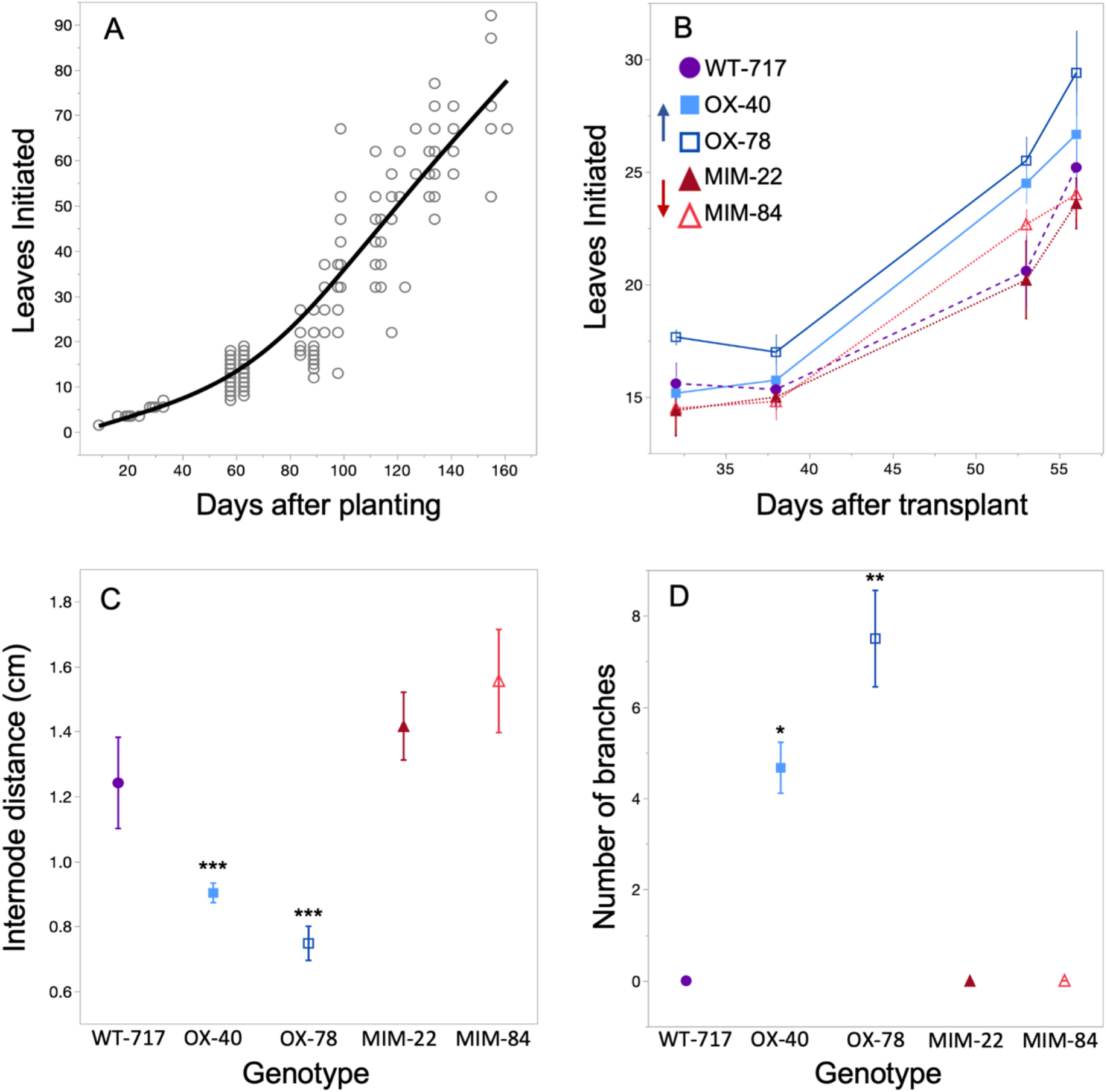
Number of leaves initiated by *P. tremula x alba* grown from seed (A) and the 717-1B4 wildtype and four transgenic lines initially propagated in tissue culture (n = 3-6) (B). The distance between nodes 10-20 (n = 6) (C), and the number of branches > 1cm in length between nodes 1 and 25 after 4 months of growth (n = 6) (D) in 717-1B4 and four transgenic lines. Data from seed-grown plants presented as individual biological replicates using grey open circles and a mean best fit line in black or as means ± s.e.m with 717 wildtype data depicted by purple circles, miR156*a* overexpression lines depicted by blue squares and MIM156 lines depicted by red triangles. Data in panel A show a second-degree polynomial fit with *p <* 0.0001 and *R*^*2*^=0.92 and an ANCOVA for panel B found *p <* 0.0001 for genotype and days after transplant but no significance for an interaction between effects. One-way ANOVA for panels C and D found *p <* 0.0001 with the results of a *Student’s t* between WT and mutants represented by * *p <* 0.05, ** *p <* 0.01, and *** *p <* 0.001.

### Juvenile and adult leaves differ in leaf shape and size

The leaves of seed-grown plants became gradually rounder (decreased L:W ratio) (Fig. **4A**) and larger (Fig. **4C**) from node 1 to node 25 and did not change significantly in shape or size after this point (Fig. **4C**, **1C**). 717-1B4 displayed a similar trend, although the leaves of this clone reached their climax shape and size a few nodes earlier than seed-grown plants (Fig. **4B, D** and **5**). MIM156 lines did not differ significantly in size from 717-1B4, but the leaves of line 84 were rounder than 717-1B4 beginning at node 15. In contrast, the leaves of miR156OX lines were significantly more elongated and smaller than wild-type leaves starting from node 15 (Student’s *t* test *p <*0.05). These results suggest that changes in leaf shape and size during shoot development in poplar are partly regulated by a decrease in level of miR156.

**Figure 4.**
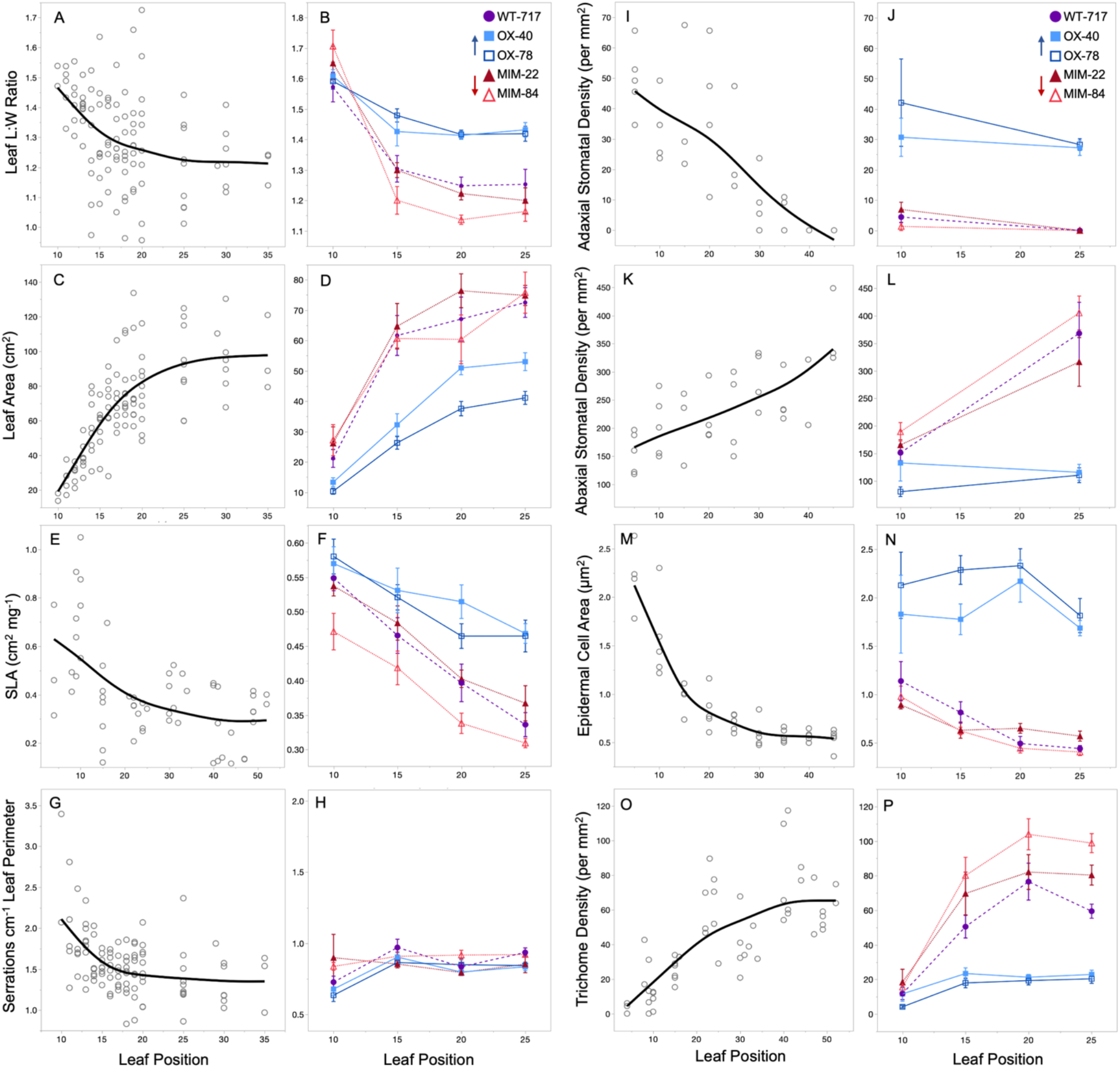
Leaf length:width (L:W) ratio, leaf area, specific leaf area (SLA), serrations per cm of leaf perimeter, adaxial stomatal density, abaxial stomatal density, adaxial epidermal cell area, and abaxial trichome density of seed-grown poplar (A, C, E, G, I, K, M, O) and the 717-1B4 and transgenic lines (B, D, F, H, J, L, N, P). Data from seed-grown plants presented as individual biological replicates using grey open circles and a mean best fit line in black or as means ± s.e.m with 717 wildtype data depicted by purple circles, miR156a overexpression lines depicted by blue squares and MIM156 lines depicted by red triangles (n = 5-11). Data in panels A, C, E, G, M and O show a second-degree polynomial relationship with *p <*0.0001 and *R*^*2*^ values of 0.17, 0.6, 0.32, 0.25, 0.85 and 0.56 respectively. Data in panels I and K show a linear relationship with *p <*0.0001 and *R*^*2*^ values of 0.58 and 0.46 respectively. ANCOVAs with leaf position and genotype as the effects for data in panels B, D, F, L, N and P found *p <*0.001 for both leaf position and genotype. In panel J, *p <*0.001 for genotype but N.S. for leaf position. In panel H, *p* < 0.01 for leaf position but N.S. for genotype. For the interaction between genotype and leaf position, *p <*0.05 for data in panels B, F, L, and P and N.S. for panels D, H, J, and N.

**Figure 5.**
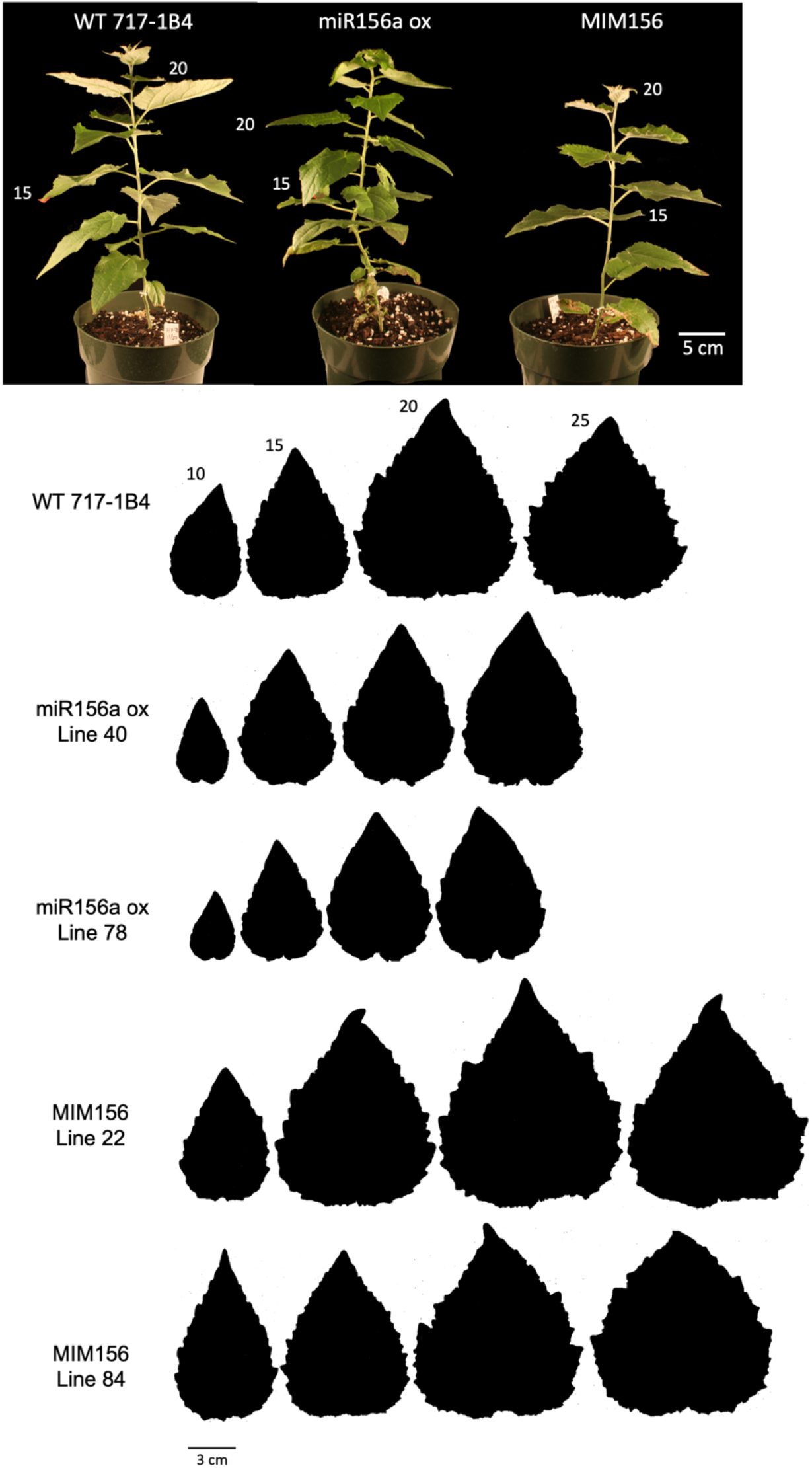
Representative plants from the wild-type 717-1B4, miR156a overexpressor line 40 and MIM156 line 84 two months following transplant from tissue culture. Leaves at positions 15 and 20 noted. Representative leaf shape for fully-expanded leaves from positions 10, 15, 20 and 25 in the 717-1B4 and transgenic poplar plants from tissue culture.

To obtain a comprehensive picture of the effect of miR156 on leaf morphology in *Populus tremula x alba*, we also measured the specific leaf area, serrations, stomatal density, epidermal cell area, abaxial trichome density, and petiole morphology of successive leaves. We examined every 5th leaf of seed-grown plants from germination to node 45-50, and every 5th leaf of 717-1B4 and transgenic lines from node 10 to 25. We observed greater variation for these traits in seed-grown plants compared to the culture-derived clones likely because of the increased genetic variation in these open-pollinated plants.

Specific leaf area (SLA) is the area of the leaf blade per unit mass, and is determined by the thickness of the leaf blade and cell density. SLA impacts leaf physiology through changes in intra-leaf light dynamics, CO_2_ diffusion and the distribution of leaf N (Parkhurst, 1994; Epron *et al*., 1995; Terashima & Hikosaka, 1995; Reich *et al*., 1998; Terashima *et al*., 2006; Evans *et al*., 2009), and therefore provides insights into how changes in leaf development alter function. SLA decreased in a curvilinear fashion from leaf 5 to leaf 40 in seed-grown plants and decreased linearly from leaf 10 to leaf 25 in 717-1B4 and the 4 transgenic lines (Figure **4E, F**). However, SLA decreased nearly linearly between leaf 10 and leaf 25 in seed-grown plants, so these data do not reveal if there is a significant difference in the pattern in seed-grown and clonally propagated plants. On the other hand, we found that—at nodes 15-25—lines over-expressing miR156 had a significantly higher SLA, and one MIM156 line (84) had a significantly lower SLA, than 717-1B4 at node 10 (Student’s *t* test, *p* <0.05 and *p* <0.0001 for miR156OX lines at nodes 15 and 20, and node 25 respectively and *p* =0.019 for line 84 node 10). These results suggest that SLA is a phase-specific trait and is regulated by miR156.

Leaf serrations are present on leaves of all measured positions for both seed- and culture-derived plants. To account for significant differences in leaf size, serration numbers are normalized to leaf perimeter. We found there to be a slight decrease in the number of serrations per cm of leaf perimeter in seed-grown plants, decreasing from just over 2 at node 10 to around 1.5 at node 35 (Fig. **4G**). 717-1B4 lines showed no phase-specific differences in the number of serrations (*p* = 0.1469) as leaves of all genotypes and all positions had between 0.64 and 0.97 serrations per cm leaf perimeter (Fig. **4H**). It is unclear as to why culture-derived plants have lower serration density than seed-grown plants however, these small differences in serrations are not likely to have any biological significance.

Stomatal density on both the adaxial and abaxial side of the leaf changed across leaf positions, but in opposite directions (Fig. **4I, K**). In seed-grown plants, adaxial stomatal density decreased linearly from leaf 5 to leaf 40, whereas abaxial stomatal density increased linearly over this interval; at every node, the density of stomata on the adaxial surface of the leaf was significantly less than that on the abaxial surface. Both 717-1B4 and the MIM156 lines had a very low density of adaxial stomata on leaf 10, and this number was reduced to nearly zero on leaf 25. In contrast, the density of abaxial stomata in these genotypes increased significantly between leaf 10 and leaf 25. In the miR156OX lines, the density of stomata on both the adaxial and abaxial surfaces of leaves 10 and 25 was essentially identical to that of the basal-most leaves of seed-grown plants. Thus, miR156 has a role in regulating stomatal density on both sides of the leaf. However, other factors are also likely to contribute to the developmental changes in stomatal density (Berger & Altmann, 2000; Sugano *et al*., 2010), because this parameter changes linearly during shoot development, rather than in the curvilinear pattern characteristic of the change in miR156 levels Fig. **4J, L**).

We measured epidermal cell area by taking impressions of the adaxial epidermis, because the high density of trichomes made it impractical to measure this trait on the abaxial surface. In seed-grown plants, cell area decreased approximately 4-fold over the first 30 leaves and remained relatively constant after this point (Fig. **4M**). In 717-1B4, cell area decreased about 2-fold from leaf 10 to leaf 20, and decreased much less dramatically, if at all, between leaf 20 and leaf 25. The MIM156 lines were not significantly different from 717-1B4. In contrast, cell area in the miR156OX did not change significantly between leaf 10 and leaf 25 (Fig. **4N**). These results suggest that epidermal cell area is a phase-specific trait.

The leaves of *Populus alba* appear white because of a dense layer of trichomes on the abaxial surface of the leaf blade, and this trait is also characteristic of *P. tremula x alba* (Fig **S3**). All the leaves of *P. tremula x alba* produced abaxial trichomes, but trichome density increased dramatically during shoot growth (Fig. **4O, P**). In seed-grown plants, abaxial trichome density increased more than 10-fold between leaf 5 and leaf 40 and increased by about 6-fold between leaf 10 and leaf 25 in 717-1B4. In contrast, abaxial trichome density remained at the same relatively low level from node 10 to node 25 in the miR156OX, and it increased significantly more than 717-1B4 in both MIM156 lines (Student’s *t* test; *p* <0.01 and *p* <0.0001 for line 22 and 84 respectively). This result suggests that abaxial trichome production is a phase-specific trait in *P. tremula x alba* and is hypersensitive to the level of miR156.

### Leaf vein density increases as plants transition from juvenile to adult

While working with the 717-1B4 lines, we noticed differences in vasculature of leaves at the same position between wildtype and transgenic lines. Because vein density and patterning impacts leaf structure and function (i.e. assimilation and herbivore resistance) (Brodribb *et al*., 2007; Blonder *et al*., 2018), we thought this trait was worth investigating. Leaf vein density increased 1.8-fold with increasing leaf position in 717-1B4 and in the MIM156 lines, although MIM156 line 84 had a significantly higher vein density than 717-1B4 in leaf 10 (Student’s *t* test, *p* <0.01). Similar results were obtained for the density of vein endings. In the miR156OX lines, vein density and the density of vein endings did not differ much between leaf 10 and leaf 25, and were identical to, or slightly lower than, the values observed for leaf 10 in 717-1B4 (Fig. **6**).

**Figure 6.**
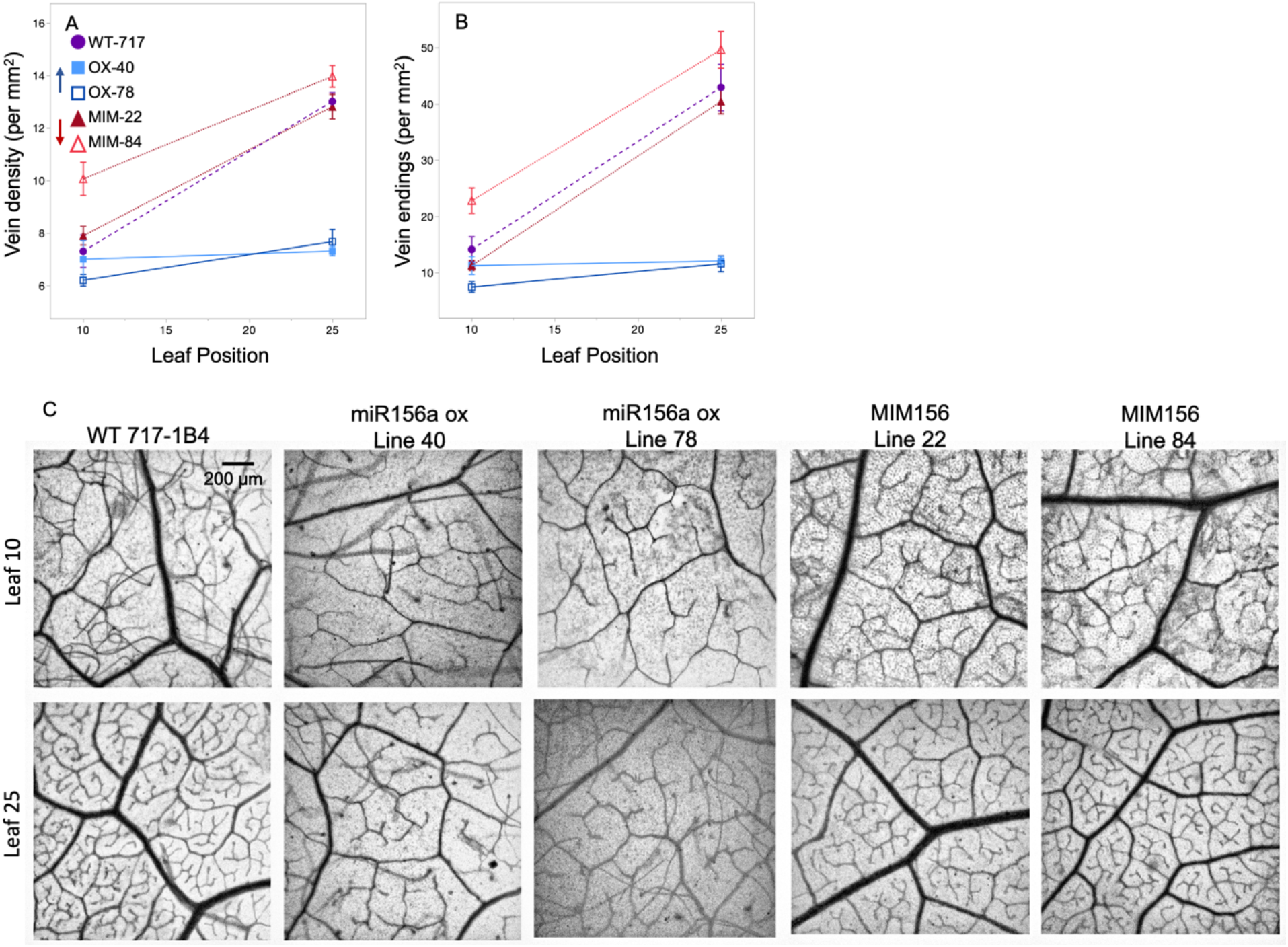
Total leaf vein density (A) and the number of vein endings (B) in the 717-1B4 and transgenic plants. Data from seed-grown plants presented as individual biological replicates using grey open circles and a mean best fit line in black or as means ± s.e.m with 717 wildtype data depicted by purple circles, miR156a overexpression lines depicted by blue squares and MIM156 lines depicted by red triangles (n = 5-6). ANCOVA with leaf position and genotype as the effects found *p <*0.0001 for both effects and the interaction for data in panels A and B. Representative samples of leaves cleared and stained to visualized vein density and architecture in leaves 10 and 25 of 717-1B4 and transgenic poplar lines (C).

### Petiole morphology changes during development leading to altered leaf fluttering behavior

Many species of poplar, including *Populus tremula x alba*, exhibit leaf fluttering, a trait that has been attributed to the vertical elongation of the petiole (Niklas, 1991). In a previous study of *Populus x canadensis*, we found that the petioles of *in vitro* propagated plants became increasing elongated as the shoot developed, and that over-expression of miR156 suppressed this process (Wang et al. 2011). To determine if this phenomenon occurs naturally, we examined the L:W ratio of a cross-section of the petiole in seed-grown *Populus tremula x alba* (Fig. **7A, D**). We found that the petiole of leaves at node 10 was slightly broader than tall (L:W < 1), whereas the petioles of leaves at higher nodes became vertically elongated (L:W >1), reaching their climax shape around node 40. In wild-type 717-1B4, and in the MIM156 and miR156OX lines, the petiole also became progressively elongated (Fig. **7B, C**). However, the miR156OX lines had significantly less elongated petioles than the 717-1B4 and the MIM156 lines, and reached a maximum L:W ratio at an earlier node. We also examined the effect of these transgenes on petiole anatomy. In 717-1B4, the petioles of leaves at nodes 10 and 15 had an adaxial indentation, a large central vascular bundle, and two smaller vascular bundles located in a horizontal plane above the central bundle. The petioles of leaves 20 and 25 lacked an adaxial indentation and had three vertically oriented vascular bundles that decreased in size from the abaxial to the adaxial side of the petiole. In the MIM156 lines, the vascular anatomy of leaf 15 was intermediate between that of leaf 10 and leaf 20 in 717-1B4, indicating that the transition to the adult shape is accelerated in these lines, despite the lack of a significant change in the L:W ratio of the petiole (Fig. **7C**). Consistent with their L:W ratio, lines over-expressing miR156 retained a juvenile petiole morphology up to at least node 25. Petiole anatomy is therefore an obvious qualitative indicator of the phase identity of a leaf in *Populus tremula x alba*.

**Figure 7.**
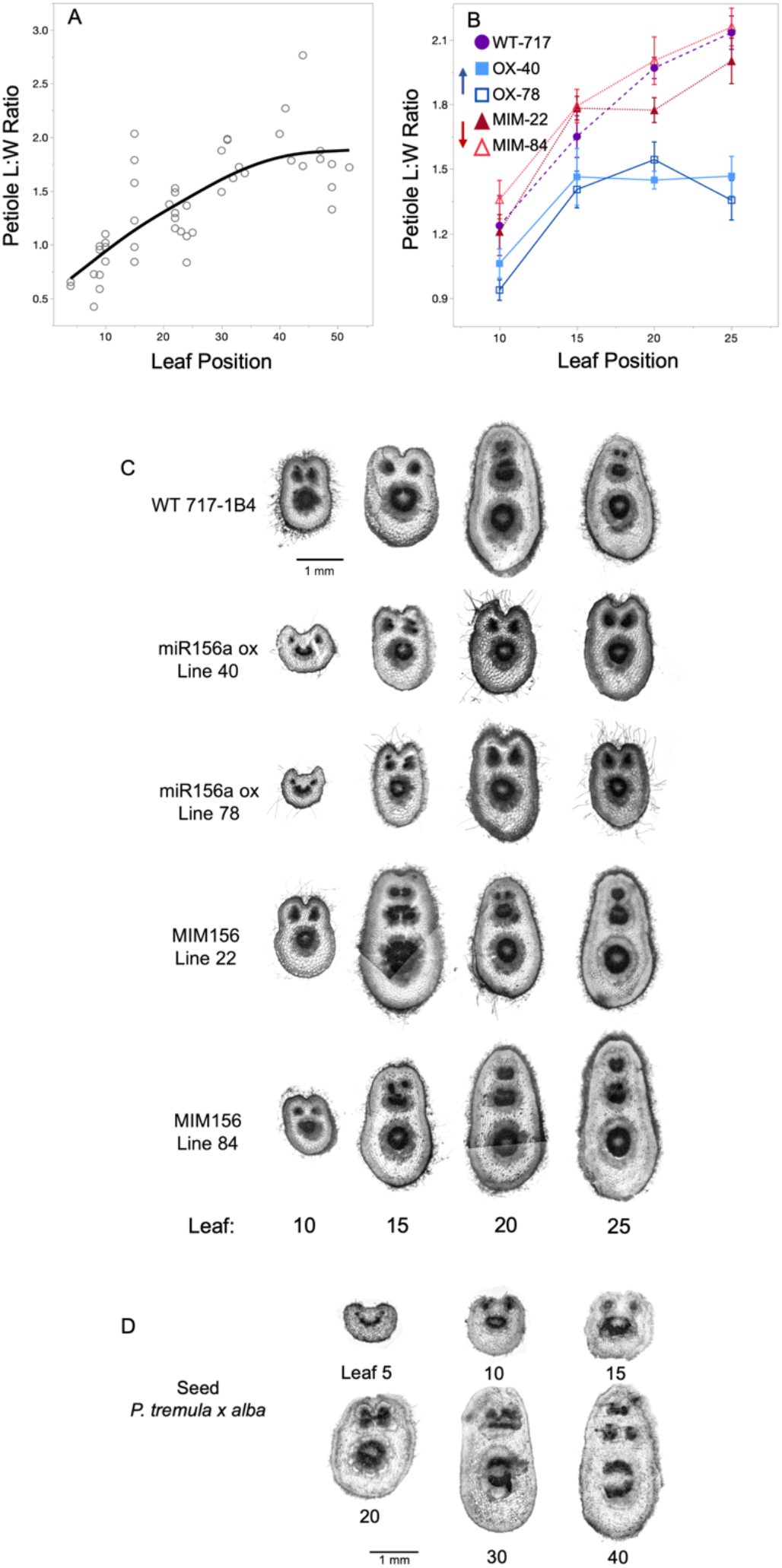
Petiole length:width ratio from cross sections made within 1 cm of leaf attachment in seed-grown (A) and 717-1B4 and transgenic poplar (B). Data from seed-grown plants presented as individual biological replicates using grey open circles and a mean best fit line in black or as means ± s.e.m with 717 wildtype data depicted by purple circles, miR156a overexpression lines depicted by blue squares and MIM156 lines depicted by red triangles (n = 7-12). Data from panel A show a second-degree polynomial relationship with *p <*0.0001 and *R*^*2*^ = 0.62. For panel B, ANCOVA with leaf position and genotype as the effects found *p<* 0.0001 for leaf position and genotype and *p=*0.01 for the interaction. Representative samples of petiole cross sections from leaves 10, 15, 20 and 25 of the 717-1B4 and four transgenic lines (C). Representative samples of petiole cross sections from leaf positions between 5 and 40 in plants grown from seed (D).

To determine if this change in petiole morphology has an effect on the behavior of the leaf, we looked at two measures of leaf fluttering—the total distance moved by the leaf, and the maximum horizontal span of the leaf tip—when leaves were exposed to a constant wind velocity (Fig. **8A**, **S4**). There was a statistically significant (*p* <0.0001) positive linear relationship between petiole L:W ratio and these fluttering measures in 717-1B4, the MIM156 and miR156OX lines. Leaves at basal node 10 in the 717-1B4 and in the MIM156 lines fluttered less than leaves at more apical node 25, whereas juvenile leaves at node 25 of the miR156OX lines fluttered less than leaves at node 25 of the 717-1B4 and in the MIM156 lines (Fig. **8B, C**). Thus, the petiole morphology of juvenile and adult leaves of *Populus tremula x alba* has a significant effect on their response to wind.

**Figure 8.**
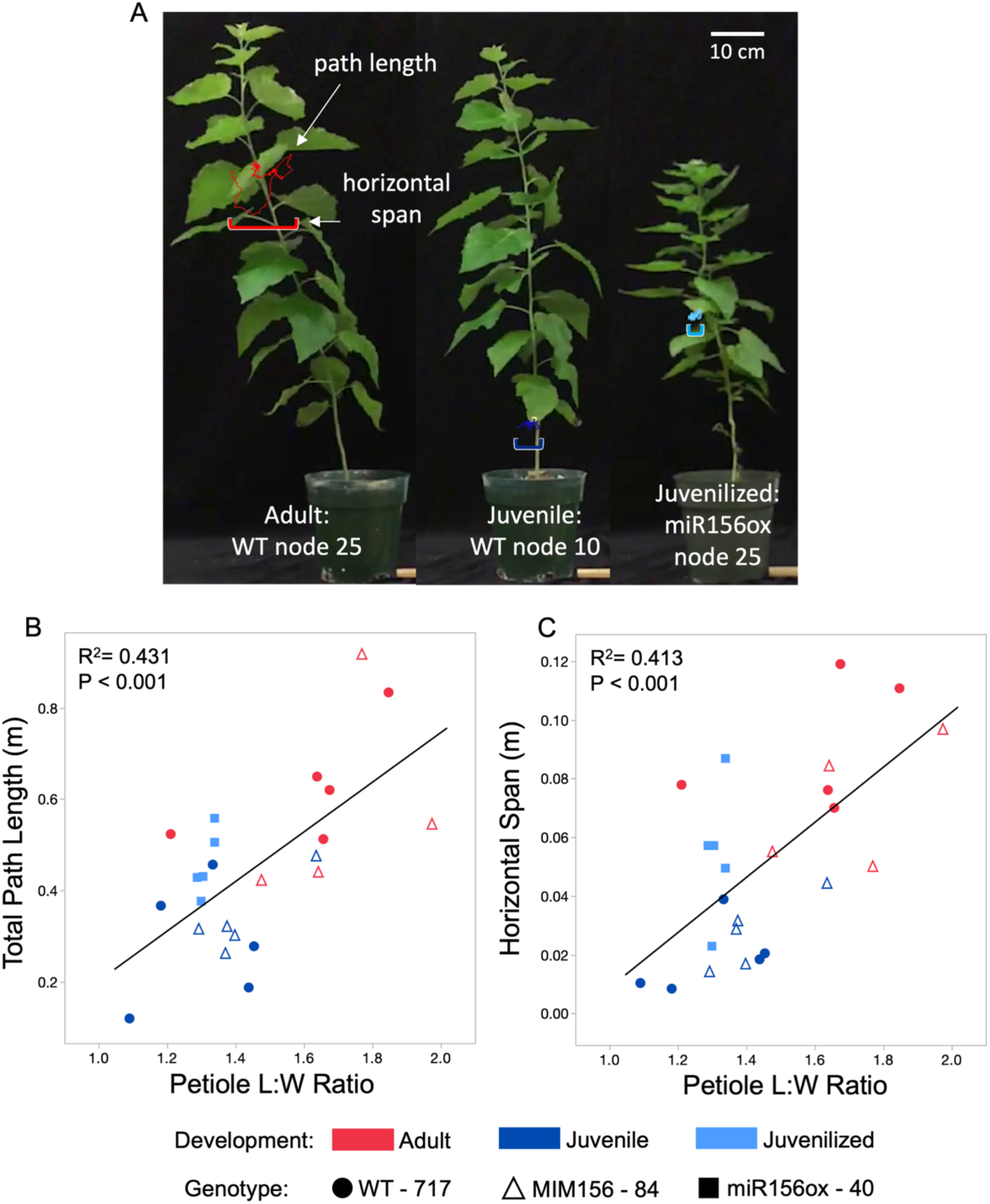
Diagram showing the path length and horizontal span measurements of leaf fluttering of an adult (WT 717-1B4 node 25), juvenile (WT 717-1B4 node 10), and juvenilized (miR156 overexpressor line 40 node 25) leaf. Quantification of leaf fluttering through total path length over a 1 second period (A) and maximum horizontal span of the tip of the leaf (B) and their relationship with petiole length:width ratio. 717-1B4 data is depicted by circles, miR156a ox line 40 by squares and MIM156 line 84 by triangles. Leaf developmental stage is noted by color with adult leaves in red, juvenile in dark blue and juvenilized in light blue. Linear fit for panel B) path length = −0.34288 + 0.544(petiole L:W ratio) and C) horizontal span = −0.0852 + 0.0939(petiole L:W ratio).

## Discussion

Many plants lack visually abrupt changes in form during shoot development, making the onset of vegetative phase change difficult to observe. As a result, the existence of this phenomenon and its significance for plant ecology and evolution are often under appreciated. In this study, we characterized the growth rate, developmental morphology, and temporal pattern of miR156 and *SPL* expression in *Populus tremula x alba*, an ecologically and economically important tree. We found that many leaf traits change during shoot development in plants grown from seeds, as well as in shoots arising *de novo* in culture, and that most of these changes are correlated with and sensitive to changes in the level of miR156. These data confirm and extend previous studies of the effect of miR156 on *Populus* development (Wang *et al*., 2011; Rubinelli *et al*., 2013), and provide a foundation for future research on the function of vegetative phase change in the life history of this woody species.

In nature, poplars propagate primarily from adventitious shoots, whereas clones used for research or commercial purposes are propagated from axillary buds or by inducing shoot regeneration *in vitro* (Taylor, 2002). The extent to which these plants recapitulate the development of the primary shoot is still unknown. We found by at least the 10^th^ node, the development of a regenerated shoot is remarkably similar to that of a seed-grown plants. This result suggests that the phase-identity of poplar cells is either re-set during the de-differentiation of the primary tissue sample, or during shoot regeneration, and is consistent with observations in *Zea mays* (Irish & Karlen, 1998; Orkwiszewski & Poethig, 2000). This observation also raises the long-standing question of whether differentiated traits can be maintained in tissue culture.

Although vegetative phase change sometimes involves traits unique to a particular species (Wang *et al*., 2011; Leichty & Poethig, 2019), analyses of plants with altered levels of miR156 have revealed a number of phase-specific traits that are common to phylogenetically diverse species. In *Zea mays, Arabidopsis thaliana, Nicotiana tabacum*, and now *Populus tremula x alba*, these traits include patterns of trichome production, leaf shape and size, epidermal and stomatal cell traits, vein density, leaf initiation rate, and branching (Bongard-Pierce *et al*., 1996; Telfer *et al*., 1997; Feng *et al*., 2016; He *et al*., 2018). At least some of these traits are also regulated by miR156 in *Populus x canadensis* (Wang *et al*., 2011), rice (Xie *et al*., 2006), alfalfa (Aung *et al*., 2015), switchgrass (Fu *et al*., 2012), *Lotus japonicus* (Wang, 2014), and tomato (Zhang *et al*., 2011). This observation indicates that vegetative phase change is an ancient and fundamental aspect of shoot development, and also suggests that the differential expression of various traits during shoot development contributes to plant fitness and is therefore under natural selection (Leichty & Poethig, 2019).

Determining how phase-specific traits contribute to plant fitness is challenging. However, a number of studies have shown that plants over-expressing miR156 are more or less sensitive to various environmental stresses (Stief *et al*., 2014; Cui *et al*., 2014; Arshad *et al*., 2017; Kang *et al*., 2020), Despite this, there is still little evidence that juvenile and adult phases of wild type plants display these same differences in sensitivity or how they might contribute to plant fitness. There are relatively few examples of phase-specific vegetative traits that are either known or expected to be selectively advantageous (Williams *et al*., 1998; Holeski *et al*., 2009; Leichty & Poethig, 2019). Our observation that adult leaves flutter more than juvenile leaves is another such example. In poplar, leaf fluttering is beneficial under high-light and high-heat environments, but is disadvantageous under low light conditions as it reduces photosynthesis (Roden & Pearcy, 1993a,b). This suggests that reduced leaf fluttering is advantageous early in development, when plants are more likely to be shaded, whereas increased fluttering is advantageous once plants are no longer in the understory.

Surprisingly, we found that progression of vegetative phase change in poplar, a woody perennial, is not very different from this process *Arabidopsis*, an annual herb. In *P. tremula x alba*, the onset of adult traits begins within three months of growth, despite the much longer lifespan of this species. As in *Arabidopsis* (He *et al*., 2018), miR156 expression in poplar declines rapidly over the first few leaves, but it isn’t until later, when the decline in miR156 is more subtle, that adult morphological traits first appear. This result further supports the findings in *Arabidopsis* that miR156/157 expression in early leaves far exceeds the amount necessary to repress most *SPL* targets (He *et al*., 2018). Further, these finding suggest that it is inappropriate to assume the timing of vegetative phase change is proportional to the lifespan of a species. As we continue to characterize this developmental transition across species with diverse life histories, we are likely to find large variations in the proportion of time spent in either the juvenile or adult vegetative phase.

During shoot development, different morphological and physiological traits change in a coordinated fashion, but not necessarily at the same node or at the same rate (Hackett & Murray, 1997; Leichty & Poethig, 2019). Furthermore, under certain conditions, traits can become dissociated, resulting in leaves or shoots that have unusual combinations of juvenile and adult traits (Robbins, 1960; Doorenbos, 1965). In our experiments, this was apparent from the phenotype of transgenic lines expressing a miR156/miR157 target site mimic (MIM156). The stronger of these transgenes had a significant effect on leaf L:W ratio, SLA, epidermal cell area, petiole morphology and leaf vein density, but did not affect leaf initiation rate, stomatal density and petiole L:W ratio. These observations are consistent with studies in *Arabidopsis*, which show that different miR156-regulated *SPL* genes have different transcription patterns and sensitivity to miR156 abundance (Xu *et al*., 2016; He *et al*., 2018). Along with the evidence that *SPL* genes also have different loss-of-function phenotypes (Xu *et al*., 2016), these results suggest that variability in the expression of different phase-specific traits may arise from the differential response of functionally distinct *SPL* genes to changes in the level of miR156.

miR156 and its *SPL* targets control many economically important traits, including flowering time, shoot architecture, biomass and lignin production, pest and disease resistance and abiotic stress responses (Wang & Wang, 2015). In the biofuel crop *P. virgatum*, for example, over-expressing miR156 leads to increases in biomass, digestibility, and lignin production, a delay in flowering time, and altered susceptibility to the pathogen *Puccinia emaculata* (Chuck *et al*., 2011; Fu *et al*., 2012; Baxter *et al*., 2018). Similar effects of miR156 over-expression have been observed in the forage legumes *Trifolium pretense* (Zheng *et al*., 2016) and *Medicago sativa* (Aung *et al*., 2015). Poplars are fast-growing trees useful for a wide variety of industrial purposes, as well for the re-forestation of degraded land (Taylor, 2002). The ability to engineer their chemistry and vegetative development through the manipulation of miR156 offers new opportunities for the improvement of this important group of plants.

## Supporting information

Leaf Fluttering Video

## Acknowledgements

We thank Samara Gray and Joshua Darfler for their assistance in caring for the plants used in this study. This research was funded by the NSF Graduate Research Fellowship (DGE-1845298), U. of Pennsylvania SAS Dissertation Research Fellowship and the Peachey Research Fund to E.H.L., NIH T32GM008216 to A.R.L and NIH GM51893 to R.S.P.

## Author Contributions

E.H.L., A.R.L., and R.S.P. planned and designed the research. E.H.L. and A.R.L. performed the experiments. C.M., and S.H.S. helped create the transgenics used in this study. E.E.D performed follow-up experiments during revision. E.H.L performed statistical analyses and wrote the first draft of the manuscript. All authors revised and provided comments on the manuscript.

## Notes

### Competing Interest Statement

The authors have declared no competing interest.

### Summary of Updates

This version of the manuscript has been revised to include more details regarding the validation of transgenic lines, gene expression at earlier developmental time points and SPL gene expression in seed-grown and culture derived plants (SPL genes are targeted by miR156, the master regulator of vegetative phase change).

