## Supplemental Figures for "Vegetative phase change in *Populus tremula x alba*"

Supplemental Material

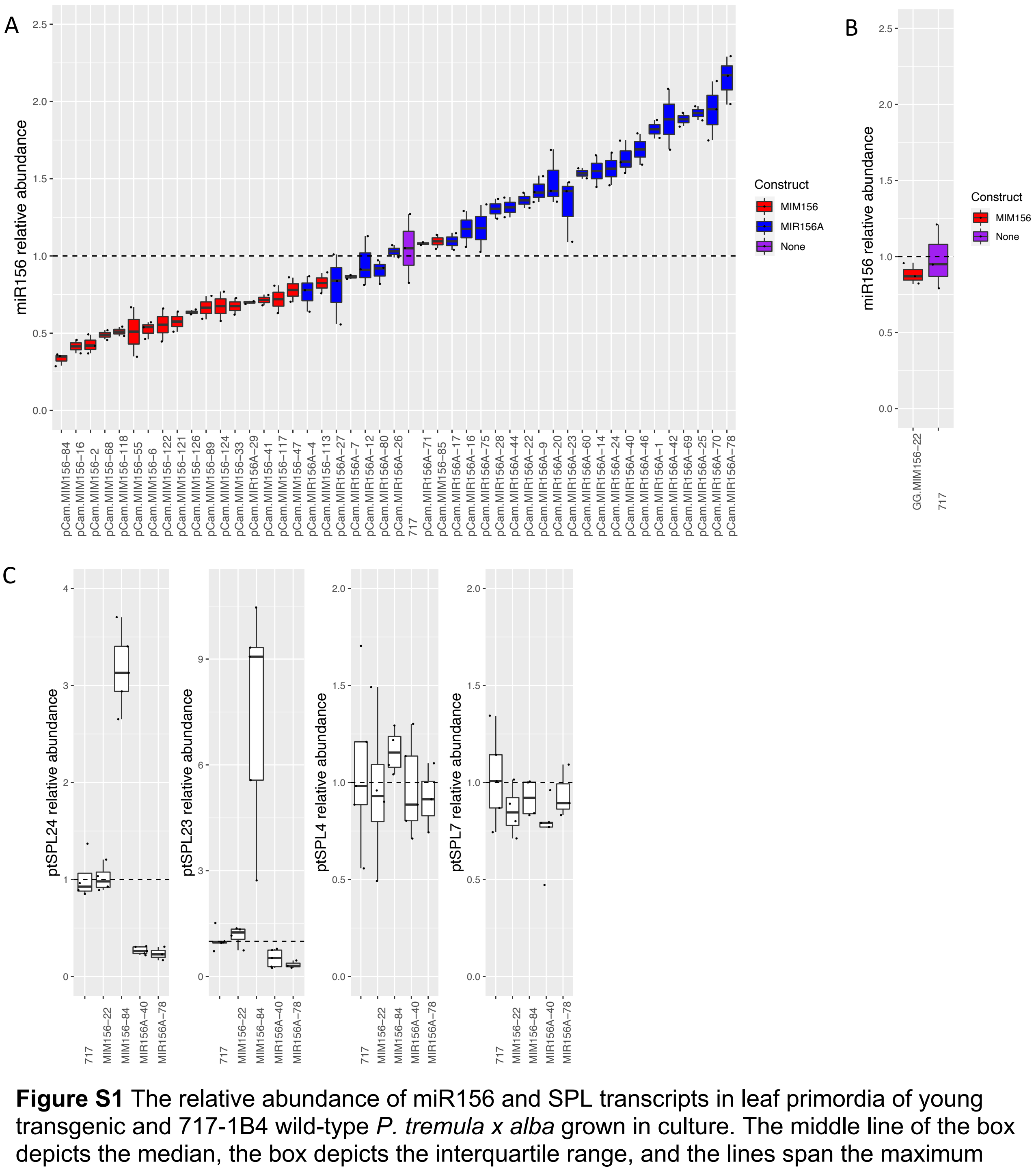


**Figure S1** The relative abundance of miR156 and SPL transcripts in leaf primordia of young transgenic and 717-1B4 wild-type *P. tremula x alba* grown in culture. The middle line of the box depicts the median, the box depicts the interquartile range, and the lines span the maximum and minimum. Individual replicates are shown as black circles. (A and B) miR156 abundance in transgenic lines over-expressing MIM156 (red) and miR156a (blue), and wild type 717-1B4 line used for transformation (purple). “pCam” = pCambia backbone, “GG” = Golden Gate backbone. ( C ) The relative abundance of SPL transcripts that contain a miR156 target site (ptSPL24 and 23), and SPL transcipts (ptSPL4 and 7) that do not.

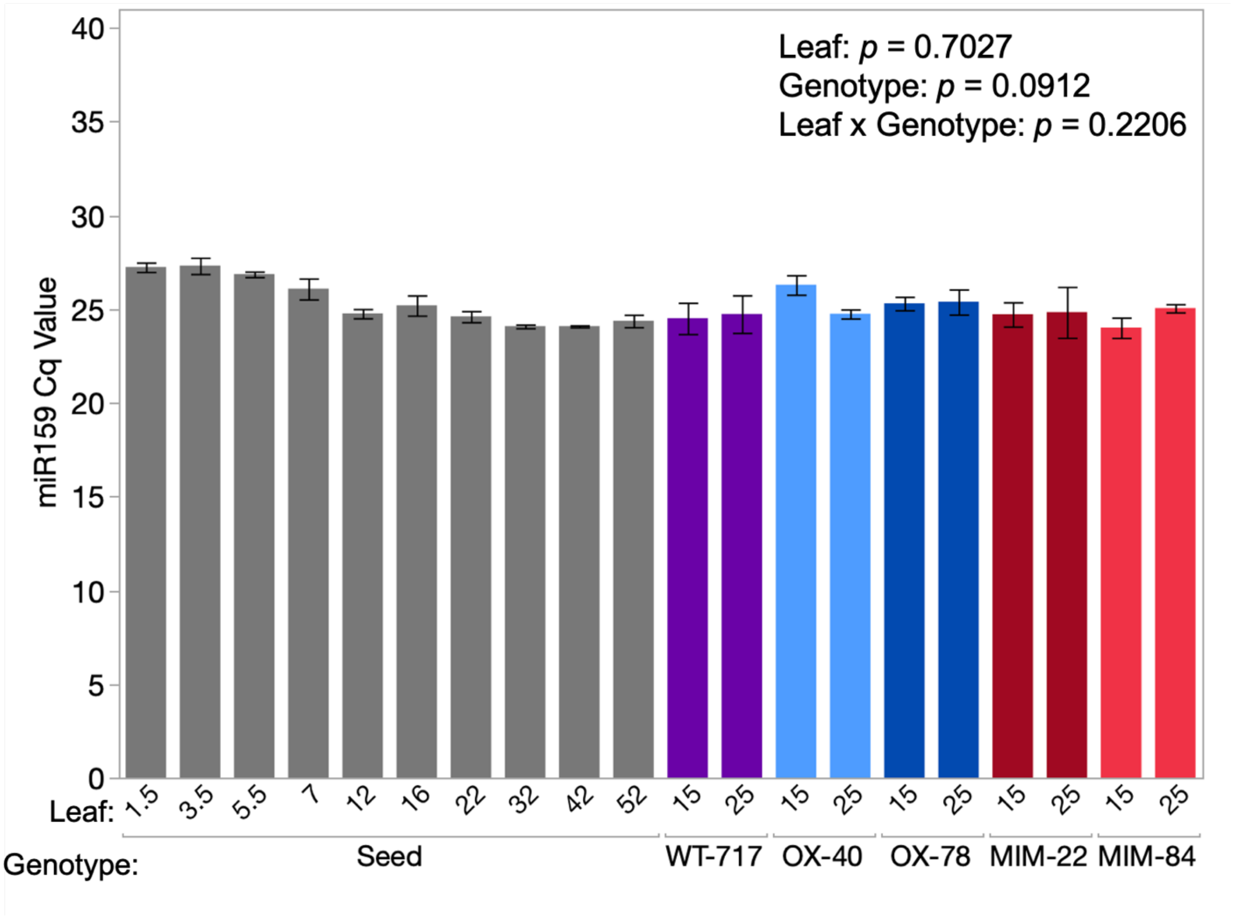


**Figure S2.** Cq values from qPCR of miR159 at different leaf positions in the genotypes of *P. tremula x alba* used in this study. These values were used to normalize miR156/7 abundance in these samples. *ANCOVA* results with leaf position and genotype as effects reported in the upper right corner of the graph show no significant differences.

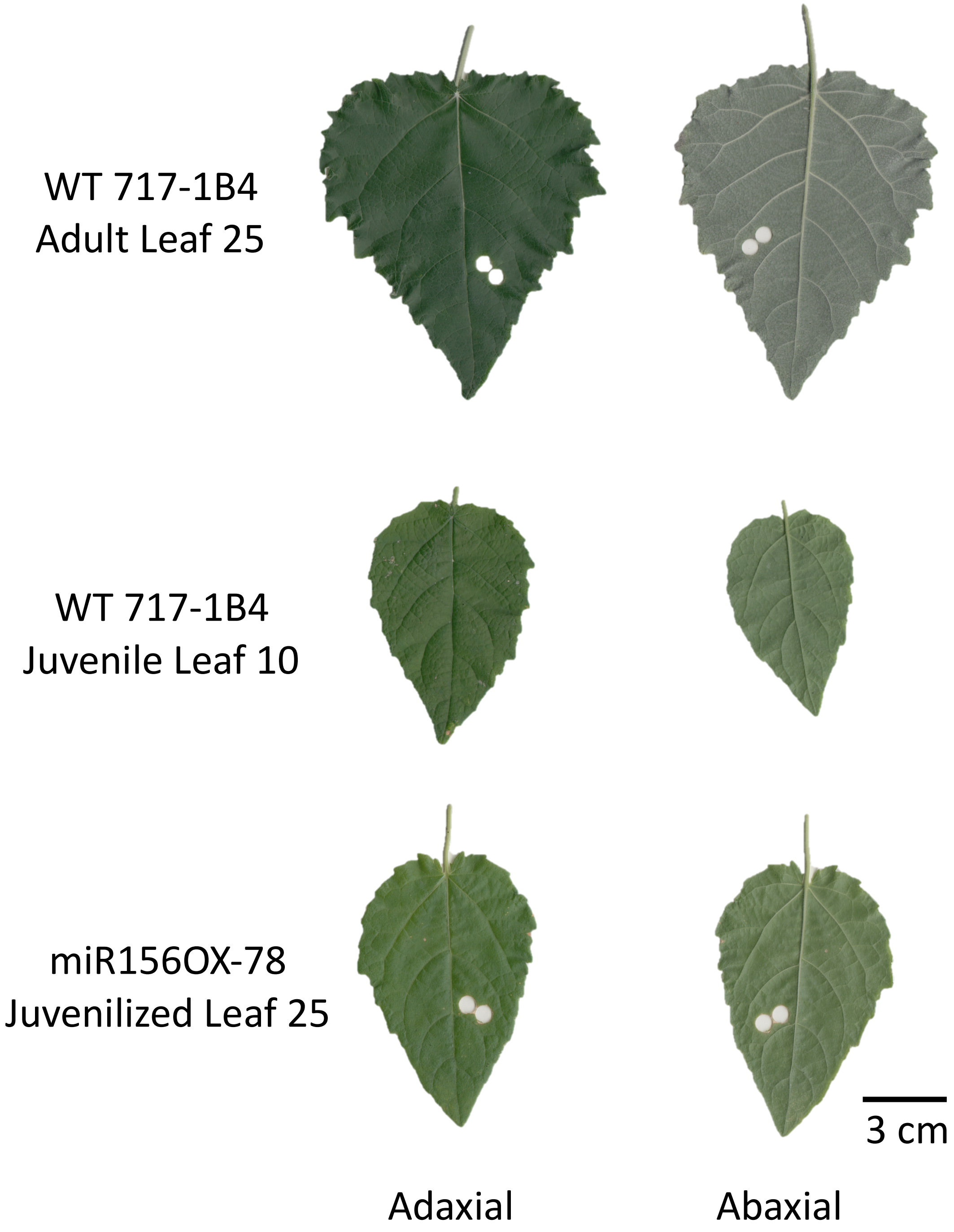


**Figure S3**. Leaf scans showing the adaxial (left) and abaxial (right) sides of adult, juvenile and juvenilized leaves of *P. tremula x alba.* Note the white appearance of the abaxial side of the adult leaf is due to a dense layer of trichomes that is not present on juvenile and juvenilized leaves. The white circles are the positions from which samples of the lamina were taken.

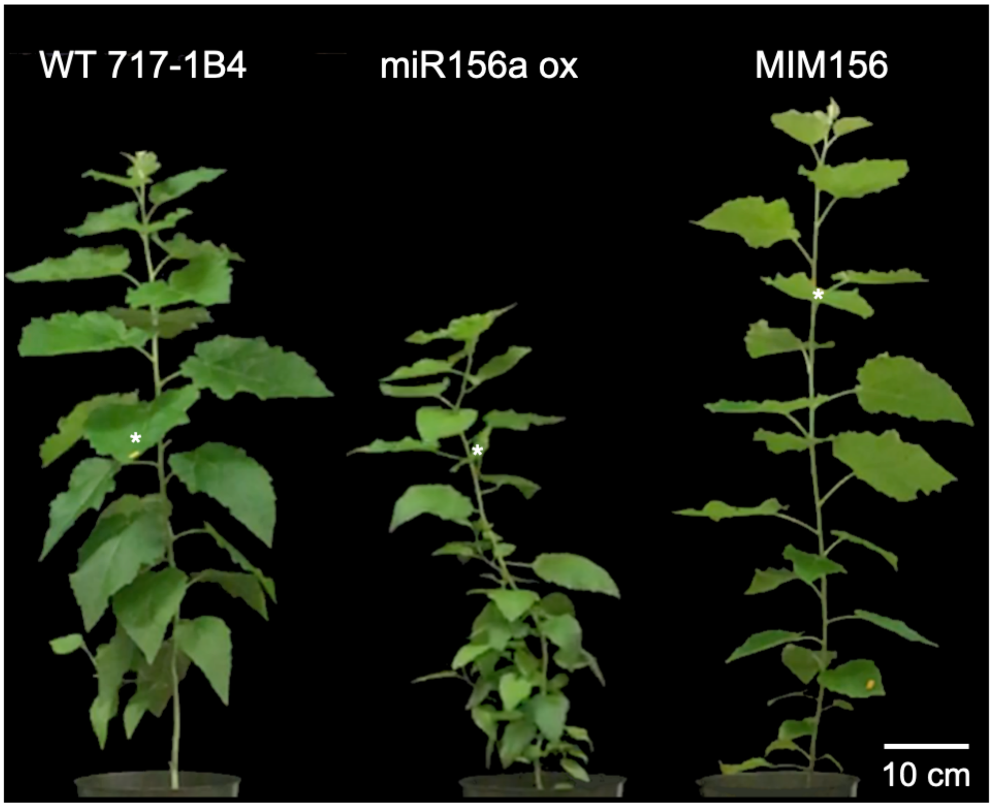


**Figure S3.** Representative plants from the wild-type 717-1B4, miR156a overexpressor line 40 and MIM156 line 84 three months following transplant from tissue culture. White asterisk marking leaves at node 25.

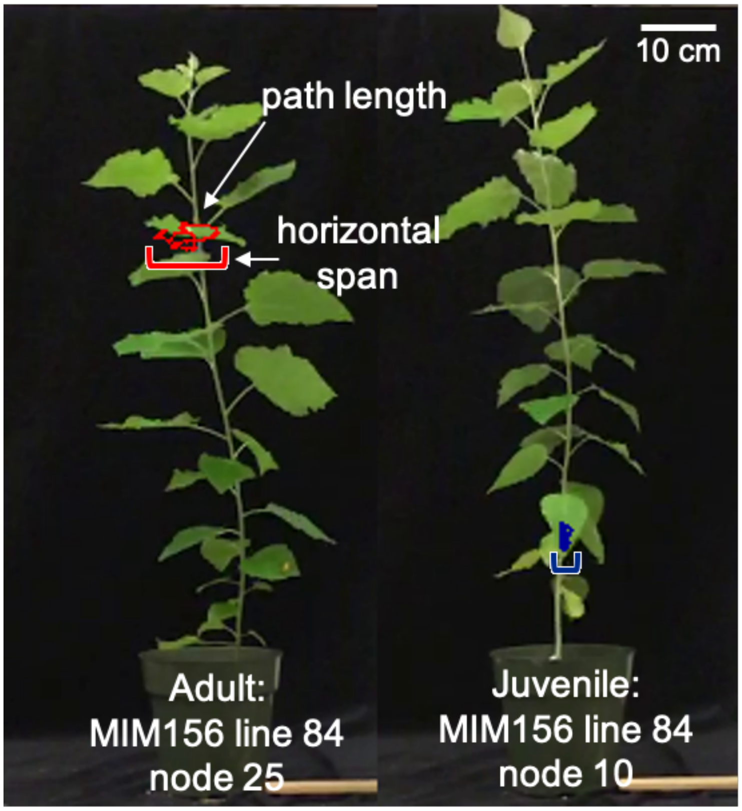


**Figure S4.** Diagram showing representative fluttering paths for adult (node 25) and juvenile (node 10) leaves of MIM156 line 84.
